# Reversibility of Nuclear and 3D Genomic Changes in Non-Cancerous Fibroblasts After Constricted Migration

**DOI:** 10.64898/2026.01.28.701973

**Authors:** Christopher Playter, Samuel John Benson, Renata Dos Reis Marques, Preston Young, Timothy B. Simmons, Kathryn Perry, Amanda K. Swets, Rachel Patton McCord

## Abstract

Metastatic cancer cells and healthy fibroblasts must traverse constrictive spaces to reach secondary sites. After passing through multiple constrictions, cancer cells often experience stable changes to their nucleus morphology, 3D genome structure, and migratory phenotype. Here, we investigate whether fibroblasts (BJ-5ta), which are non-cancerous and have an inherent ability to migrate to fulfill roles in wound repair, likewise experience nuclear and 3D genomic changes with constricted migration. We find that BJ-5ta cells do not get progressively better at migrating with sequential attempts but do experience nuclear deformations and 3D genome alterations at the compartment level after constricted migration. Transient compartment shifts spatially rearranged genes associated with preparation for and response to migration. Unlike the stable changes associated with long term phenotype changes in cancer cells, however, the nucleus deformations recovered back to unmigrated levels following proliferation and cell movement. Some compartment changes persist and might influence responses to future stimuli, but most 3D genome changes revert to the unmigrated state after cell proliferation. Our study shows that non-cancerous migratory cells are not more robust against alterations caused by constricted migration but can recover from the ones that do arise more readily than cancer cells.

**Significance Statement:**

- Cancer cells experience stable phenotype shifts and 3D genome changes after constricted migration, but it is unknown whether these changes would occur in innately migratory non-cancerous fibroblasts.
- Fibroblasts (BJ-5ta) show reversible nuclear morphology and 3D genome alterations after constricted migration.
- Our results reveal the connection between basic nucleus mechanobiology and genome structure in fibroblasts, showing how these cells adapt and respond to the forces of constriction during processes such as wound healing.

## Introduction

The ability to pass through constrictive pores is a hallmark of metastatic cancer cells, as they move from primary to secondary sites of proliferation (Fidler, 2003; Gupta and Massague, 2006; Lambert *et al*., 2017; Golloshi *et al*., 2022), but is also key to the healthy functions of non-cancerous cell types such as fibroblasts and neutrophils (Singer and Clark, 1999). For cells to pass through these constrictions, they must squeeze their nucleus, a large and stiff organelle, exerting substantial forces on its envelope and the genomic contents within (Friedl *et al*., 2011; Fu *et al*., 2012; Davidson *et al*., 2014).

Several studies have shown, across numerous cancer cell types (including leukemia, fibrosarcoma, osteosarcoma, melanoma, and breast cancer) that this deformation of the nucleus during repeated constricted migration leads to altered cell and nucleus phenotype, chromatin state, and gene expression (Irianto *et al*., 2017; Golloshi *et al*., 2022; Hsia *et al*., 2022; de Lope-Planelles *et al*., 2023). In our previous work on human melanoma A375 cells, for example, we observed that 10 rounds of constricted migration resulted in cells with stably increased migratory potential, nuclear deformations, gene expression alterations, and changes in their 3D spatial compartmentalization of heterochromatin and euchromatin (Golloshi *et al*., 2022; Playter *et al*., 2025). However, it is less apparent whether such changes would be caused by constricted migration in non-cancerous cells. Previous work showed far less dramatic 3D genome structure changes in neutrophils after constricted migration (Jacobson *et al*., 2018). But, neutrophil nuclei are inherently more deformable as they do not express the Lamin A/C protein that contributes to the rigidity of nuclei in other migratory non-cancerous cells such as fibroblasts (Rowat *et al*., 2013; Harada *et al*., 2014; Stephens *et al*., 2017b). Further, neutrophils do not survive long enough to enable testing the effect of sequential constricted migrations or the long-term stability of any resulting phenotypic changes. Therefore, in this work, we investigate the impact of single or sequential constricted migration on the phenotype, nucleus structure, and 3D genome structure of non-cancerous human fibroblasts.

Fibroblasts are innately migratory, having to traverse long distances in response to chemical cues to reach sites of wound repair, and must traverse small pores in the dense extracellular matrix (ECM) during this migration (Singer and Clark, 1999). Previous studies on fibroblast migration in 2D (scratch assays) (Li *et al*., 2004; Acharya *et al*., 2008; Hade *et al*., 2024) and in 3D (Rhee and Grinnell, 2007; Carlson *et al*., 2008) have revealed features of fibroblast migration, including the importance of the interplay between mechanical properties of the cellular environment and of the nucleus. Nucleus movement and positioning are deliberately controlled during fibroblast migration. Before forward migration of the cell, the nucleus is actively moved toward the rear of the cell by transmembrane actin-associated nuclear (TAN) lines anchored to Lamin A/C(Chang *et al*., 2013), and nucleus rotation during fibroblast movement on 2D surfaces is constrained by the focal adhesion-anchored actin cap connected to the nucleus through LINC complexes(Luxton *et al*., 2010; Zhu *et al*., 2018). Fibroblasts can also use the nucleus as a piston to exert force during constricted migration through elastic 3D matrices (Petrie *et al*., 2014).

In line with the fact that the nuclear lamina and the chromatin structure are the two main components determining the physical properties of the cell nucleus (Stephens *et al*., 2017a), alterations to these factors have been shown to cause altered fibroblast migration. Altering chromatin through epigenetic modifying drugs alters fibroblast migration and invasion in the context of keloid-forming fibroblasts (Almier *et al*., 2025). Decreased expression of Lamin A/C facilitates migration of mouse embryonic fibroblasts (MEFs) through narrow channels (Davidson *et al*., 2014). But, Lamin A/C expression is actually helpful when fibroblasts migrate through dense fibrous environments, enabling the fibroblast nuclei to deform with deep invaginations around obstacles (Katiyar *et al*., 2022).

While all this evidence indicates that the nucleus experiences physical forces during constricted migration, and the physical properties of the fibroblast nucleus can affect how well it can pass through constrictions, it is not clear whether sequential constrictions would lead to stable changes in fibroblasts as have been observed in cancer cell types. A suggestion that phenotype differences could result from mechanical cues comes from the observation that fibroblasts taken from patients with fibrosis were more migratory than those from healthy tissues. The denser the fibrosis, the more migratory the fibroblasts (Suganuma *et al*., 1995), which could reflect the impact of the dense, stiff fibrotic tissue, on fibroblast migration phenotype (Chang *et al*., 2025). There is also evidence of both reversible and non-reversible changes in fibroblasts during their physiological roles in tissues. In response to stimuli in vivo, fibroblasts can be activated and become more migratory. Some fibroblasts quickly recover from this activation while others become persistently activated (Chang *et al*., 2025). Here, we investigate the question of whether the stresses of sequential constricted migration lead to an altered phenotype, nucleus structure, and chromatin structure, and whether any observed changes are reversible or persistently different.

To explore this question, we allowed BJ-5ta cells to migrate through multiple sequential rounds of constricted migrations to assess migration ability, nuclear lamina deformation, and chromatin architecture. Here, we found that BJ-5ta cells do not get progressively better at constricted migration but do experience some nuclear morphology and 3D genome alterations. Unlike previous observations in cancer constricted migration, however, these changes do not get progressively more dramatic. Instead, nucleus and lamin deformations become less dramatic over sequential rounds of migration. By imaging LaminA/C tagged with GFP in live cells, we found that nuclear wrinkles present after migration can smooth out during further cellular movement. With Hi-C data, we found that many spatial compartment alterations in the 3D genome also show the ability to recover after proliferation, though some compartment alterations do not recover. Our results suggest that while BJ-5ta cells do experience constricted migration induced changes to nuclear morphology and chromatin structures, these changes are largely reversible and do not lead to long term alterations to cell phenotype.

## Results

### BJ-5ta cells experience nuclear deformations following constricted migrations but do not get progressively better at migration over multiple rounds

To evaluate the effects of constricted migration on BJ-5ta fibroblasts, we first established a Transwell migration assay that would result in reproducible migration of these non-cancerous cells **(Figure 1A**, Methods). Unlike invasive cancer cell lines that migrate well in a simple serum gradient, the BJ-5ta cells required multiple extra steps such as overnight attachment and starvation followed by spike in of a chemoattractant, Fibroblast Growth Factor (FGF), to induce migration through the 5 µm pores. During the time course of 5 rounds of constricted migration, we found that BJ-5ta cells that have migrated through successive rounds do not show significant improvement in migration **(Figure 1B)**. The migration rate through the Transwells remained ∼30% from initial migration through 5 total rounds, while A375 melanoma cells typically increase from 20% to 60% migration over 5 rounds of constrictions (Golloshi *et al*., 2022).

**Figure 1.**
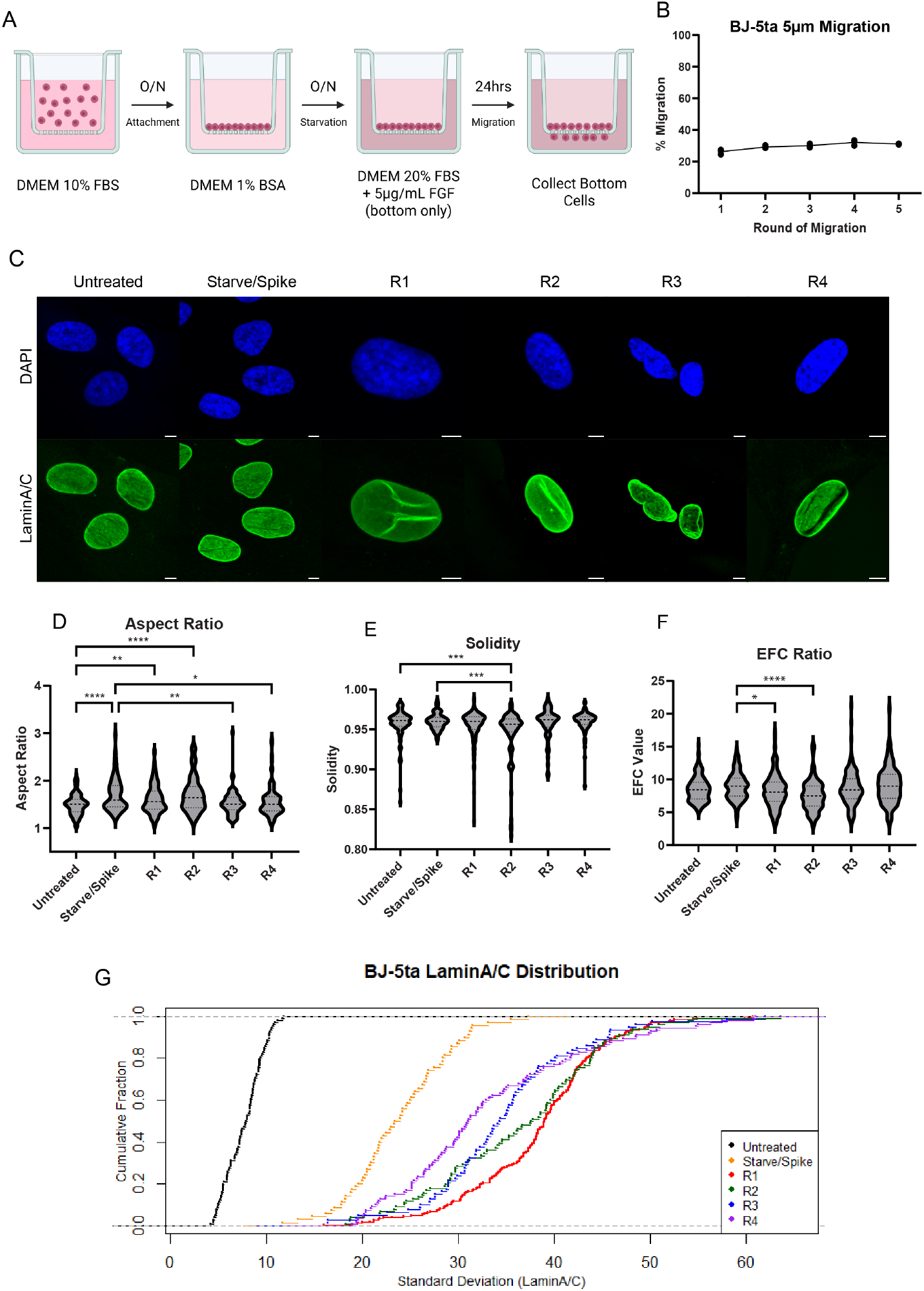
Constricted migration causes nuclear deformation and LaminA/C alterations in BJ-5ta cells. **A)** Schematic of BJ-5ta Transwell migration assay. **B)** Quantification of migration rates (% of cells that migrated through pores) through 5 µm pores for BJ-5ta cells across 5 rounds of sequential migration. **C)** Confocal images (63x) of BJ-5ta nuclei stained with LaminA/C (green) and DAPI (blue) for both unmigrated conditions and sequential migration rounds (R1-R4). Scalebar = 10 µm. Aspect Ratio **(D)**, Solidity **(E)** and EFC Ratio **(F)** measurements for conditions shown in **1C**. (****P<.0001, ***P<.001, **P<.01, *P<.05, one-way ANOVA. N = 60-250 nuclei per condition.) **G)** Cumulative distribution plot of LaminA/C standard deviation for conditions shown in **1C**. N = 60-250 nuclei per condition. In all violin plots, thick dashed lines indicate median, and thin dashes indicate 25^th^ and 75^th^ percentiles, respectively.

However, over multiple rounds of constriction we found that these fibroblasts do experience nuclear morphology differences **(Figure 1C)**. We quantified nucleus morphology using Lamin A/C immunofluorescence staining. Treating the cells with the conditions required to prime migration (starvation followed by a shift to media containing 20% FBS and 5 μg/mL FGF, which we call “starve/spike” or S/S) was able to induce morphology changes such as an increased nuclear aspect ratio, even before the cells went through migration **(Figure 1D)**. We observed that after the first 2 rounds of migration, the nucleus aspect ratio remained higher than in untreated/unmigrated (UU) cells, but was not significantly different from S/S. Interestingly, as cells passed through the 3^rd^ and 4^th^ rounds of migration, their nucleus began to return to a more rounded shape, similar to UU. This may indicate that as these cells progressed through multiple rounds of constriction, the nucleus was more adapted to handle the forces and recovered its shape more quickly after migration. In fact, when looking at nuclear solidity, a measurement of nucleus roundness, we found a significant drop in solidity relative to both UU and S/S after the 2^nd^ round of migration, but this roundness returned to unmigrated levels in rounds 3 and 4 of migration **(Figure 1E)**. The solidity measurement can be limited in its ability to detect differences due to its small dynamic range, so we also employed the Elliptic Fourier Coefficient (EFC) ratio (Tamashunas *et al*., 2020a). The smaller the value, the more irregular the nucleus. We found that after the 1^st^ and 2^nd^ rounds of constricted migration, the nuclei were significantly less regular when compared to S/S **(Figure 1F)**. Similar to what we saw with solidity, the EFC ratios returned back to levels equal to that of both unmigrated conditions as the cells passed through the 3^rd^ and 4^th^ rounds of migration. These data suggest that over rounds of constriction, BJ-5ta nucleus morphology becomes more adaptable rather than more stably distorted.

Along with nucleus shape changes, we found alterations in the distribution of lamin proteins after rounds of constricted migration. We quantified wrinkles in the nuclear lamina using the standard deviation of the Lamin A/C immunofluorescence signal for each nucleus for each condition **(Figure 1G)**. Conditions that lie further to the right in the cumulative distribution plot have larger standard deviations, usually caused by bright lines of wrinkles in the lamina. The initial starve/spike condition increases lamin wrinkling compared to untreated cells, and then the initial round of constricted migration causes the most dramatic further increase in lamin wrinkles. As cells went through more rounds of migration, the lamin standard deviation decreased sequentially as R2 was less than R1, R3 less than R2, and R4 less than R3, but still elevated compared to S/S. Combining the Lamin A/C data with the nucleus shape data, our data suggest that BJ-5ta cells experience nucleus deformations as the cells prepare for and then undergo constricted migration, but these cells can adapt to the stress of constricted migration and lessen the consequences associated with it over sequential rounds.

### BJ-5ta nuclei recover from constricted migration induced nuclear morphology changes over time

We next looked to see if the nuclear morphology changes that occurred following early rounds of constricted migration were stable in those cell populations over time. We took cells that had been migrated 1, 2, or 3 times through 5 µm pores and allowed them to proliferate for 4 days, imaging their nuclear lamina after each day of growth **(Figure 2A)**. Here, we focus on round two of migration, since it showed the largest initial changes in morphology compared to unmigrated. We found that the nucleus aspect ratio increased immediately after migration (No Growth) and then showed a downward trend over the 4 days of growth. By days 3 and 4, the aspect ratio had returned to similar levels as the unmigrated condition **(Figure 2B)**. Likewise, there was a significant reduction in both solidity and EFC ratio immediately following migration that returned to unmigrated levels over the days of growth **(Figure 2C and D)**. Solidity measurements showed that even after as little as 1 day of growth, nuclei regularity significantly increased. This recovery became even more evident over days 2-4 of growth so that the 4^th^ day of growth solidity measures were about equal to that of unmigrated cells **(Figure 2C)**. The EFC ratios showed a similar trend towards recovery after each day of growth, but statistically significant recovery was only observed following 4 days of growth **(Figure 2D)**. We saw similar trends of recovery after one round **(Sup. Figure 1A-D)** and three rounds of migration **(Sup. Figure 1E-H)**. The average doubling time for BJ-5ta cells is ∼36 hours, leading us to hypothesize that these deformations might recover as a result of cell division, where the nuclear envelope is broken down and then reassembled.

**Figure 2.**
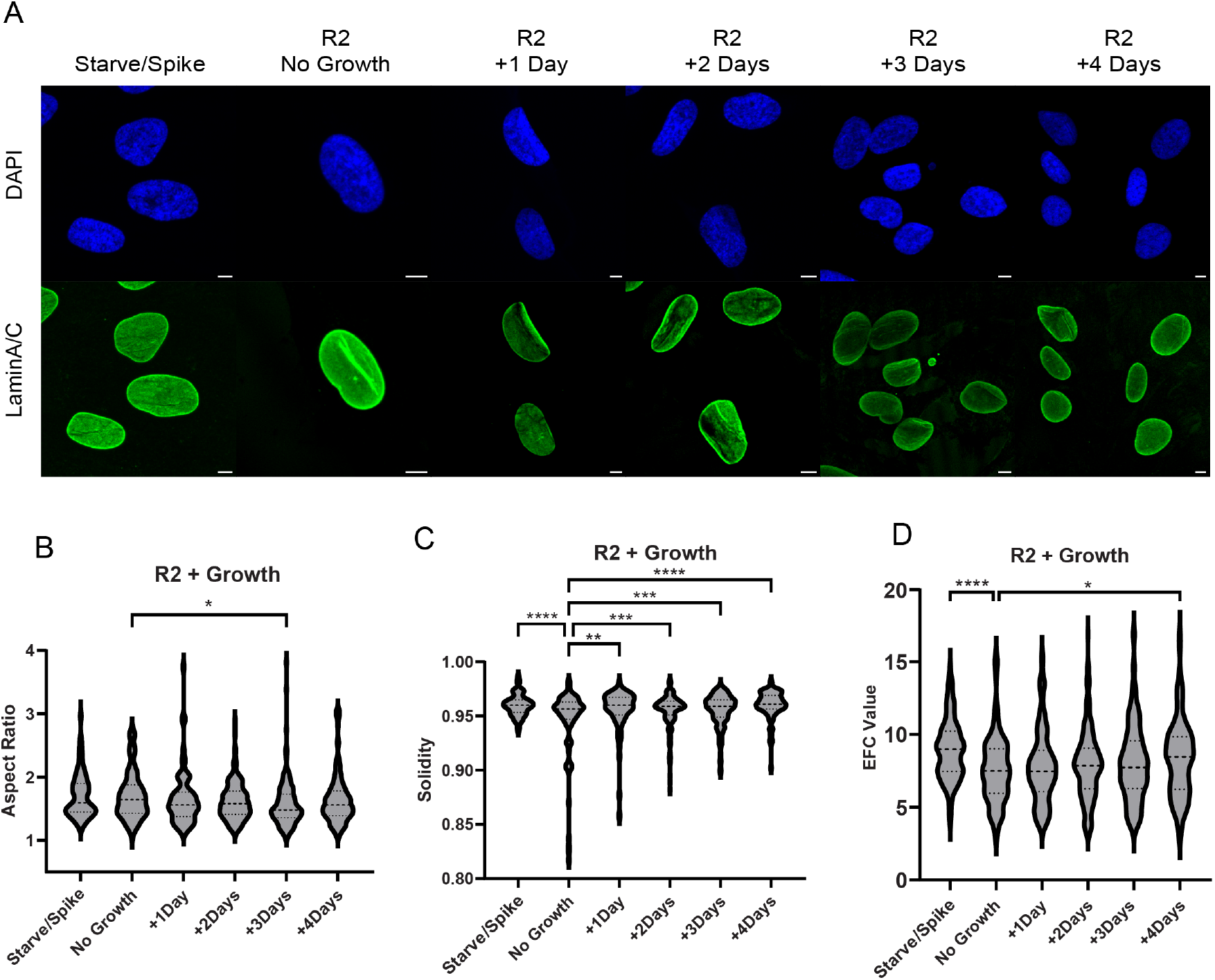
Nucleus deformation recovers over time with proliferation. **A)** Confocal images (63x) of BJ-5ta nuclei stained with Lamin A/C (green) and DAPI (blue) of S/S and R2 immediately after migration (no growth) and then followed through 4 days of proliferation. Scalebar = 10 µm. Aspect Ratio **(B)**, Solidity **(C)** and EFC Ratio **(D)** measurements for conditions shown in **2A**. (****P<.0001, ***P<.001, **P<.01, *P<.05, one-way ANOVA. N = 68-204 nuclei per condition.) In all violin plots, thick dashed lines indicate median, and thin dashes indicate 25^th^ and 75^th^ percentiles, respectively.

### Stable and dynamic nucleus deformations during 2D cell movement following constriction

Visualizing nucleus morphology only in fixed images at certain timepoints does not reveal the dynamics of nucleus shape in individual cells. We next asked whether the lamin deformations we observed were stable, requiring cell division for recovery, or whether they were dynamic within one interphase. To assess how nucleus shape and lamin wrinkles change over time, we stably transduced the BJ5-ta cells with a GFP tagged LaminA/C (GFP-LMNA), allowing visualization of LaminA/C in live cells **(Sup. Figure 2A)**. To ensure that the expression level of GFP-LMNA was biologically relevant and not too highly overexpressed, we performed fluorescence activated cell sorting (FACS) to select cells with detectable but low expression of the GFP construct **(Sup. Figure 2B and 2A white arrows)**. We then migrated these cells through 5 µm pores and visualized GFP-LMNA over time after migration. We found that some nuclei transitioned from highly wrinkled to smooth during normal cell movement without going through a division process **(Figure 3A, Movies 1-3)**. In multiple fields we saw nuclei with wrinkles that were more stable (red triangle, Nucleus 1) as well as cells that unfolded wrinkles over short amounts of time (purple circle, Nucleus 2). When tracking nucleus solidity of these two nuclei, we found that Nucleus 2 lost its wrinkles leading to an increase in solidity and a decrease in lamin standard deviation over time **(Figure 3B, C)**. Nucleus 1, in contrast, showed stable lamin wrinkles throughout the course of imaging **(Figure 3B, C)**.

**Figure 3.**
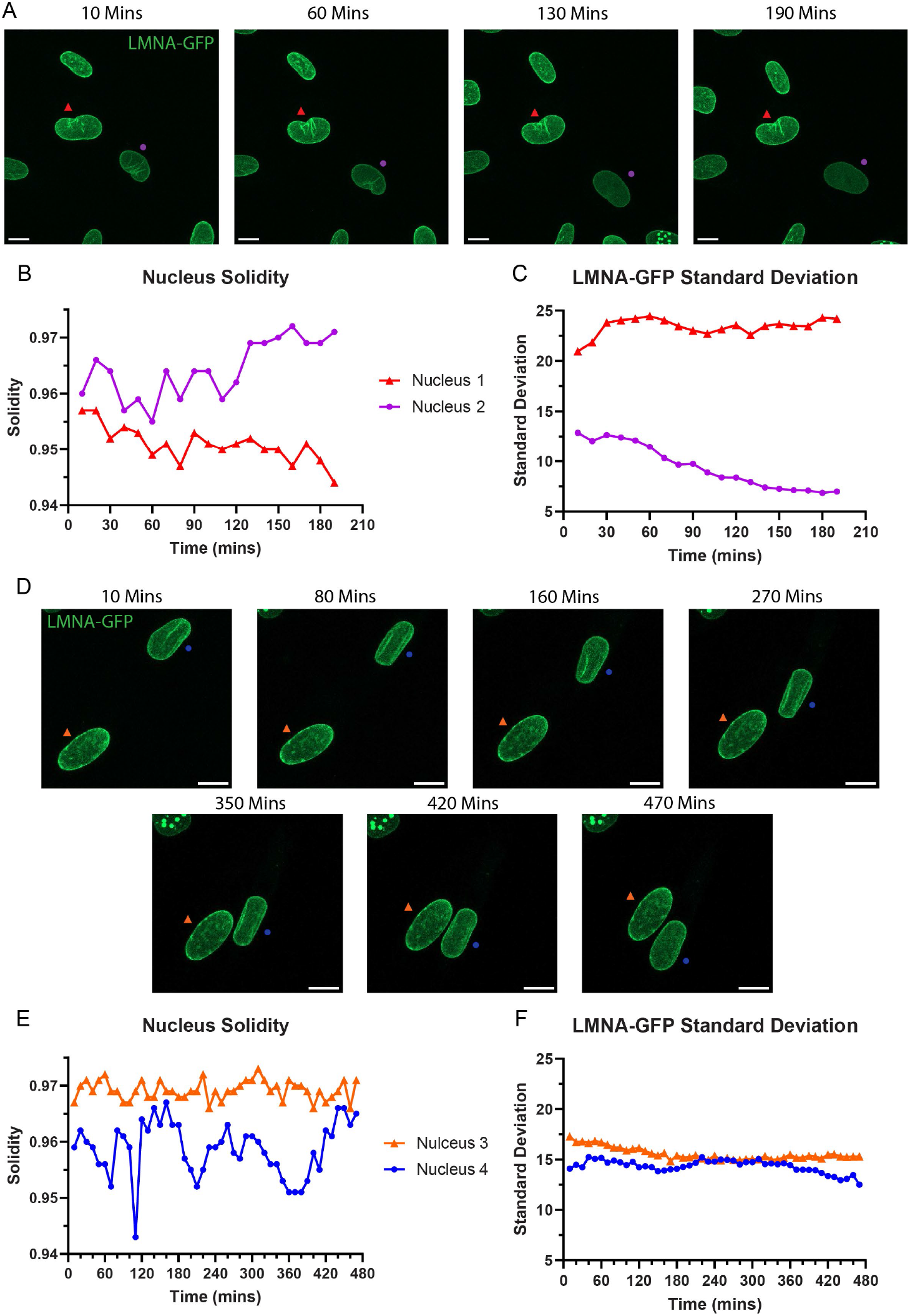
Nucleus deformations can be stable or dynamic with spontaneous cell motion. **A)** Confocal snapshots from **Movie 1** taken as cells expressing GFP-LMNA (green) move over time. Red triangle = Nucleus 1 (stable wrinkles), Purple circle = Nucleus 2 (dynamic wrinkles). Scalebar = 10 µm **B)** Solidity measurements of Nuclei 1 and 2 (from **3A**) taken at each 10min interval. **C)** GFP-LaminA standard deviation measurements of Nuclei 1 and 2 taken at each 10 min interval. **D)** Confocal snapshots from **Movie 2** taken as cells expressing GFP-LMNA move over time. Orange triangle = Nucleus 3 (stable), Blue circle = Nucleus 4 (dynamic wrinkles) Scalebar = 10 µm **E)** Solidity measurements of Nuclei 3 and 4 taken at each 10min interval **F)** GFP-LMNA standard deviation measurements of Nuclei 3 and 4 taken at each 10min interval.

We observed that lamin structure does not only progress from more to less wrinkled over time, but instead some nuclei lose and then regain lamina wrinkles as they move **(Figure 3D)**. In an example field, one nucleus (orange triangle, Nucleus 3) remains fairly static in lamin structure and position while the other (blue circle, Nucleus 4) gains and loses lamin wrinkles as it moves across the field of view. This appearance and removal of wrinkles can be seen in the fluctuation of the coinciding solidity measurements and an increase then decrease in LMNA standard deviation for Nucleus 4 **(Figure 3E, F)**. This live imaging shows that some nucleus deformations we see in fixed images are dynamic, consistent with the forces exerted on the nucleus by the cytoskeleton during cell movement.

We next tracked nuclei across cell division to determine whether the more stable lamin wrinkles would persist after division or would be lost during the nuclear envelope breakdown and reformation that happens during mitosis. We observed that cells that express GFP-LMNA display bright puncta of protein directly after cell division that diffuse gradually over time **(Sup. Figure 2C, Movie 4)**. Puncta of GFP-LMNA have been seen before in other cell lines (Hübner *et al*., 2006), and may be a property of GFP-tagged lamin. By quantifying GFP-LMNA standard deviation, we were able to determine that the puncta that arise right after division lessen in intensity over time **(Sup. Figure 2D)**. We saw similar dynamics in multiple fields where GFP-LMNA puncta appeared after division **(Movie 5 and 6)**. During a 72-hr movie, we tracked numerous dividing cells and were able to see that every dividing nucleus presented with puncta, and the puncta dissipated over the course of 15-20 hours after division, about half the length of a cell cycle for these fibroblasts **(Sup. Figure 2E, Movie 5)**. Though the formation of puncta may reflect differences between GFP-tagged and WT LaminA/C (Odell and Lammerding, 2024), these results show that dramatically different lamin structures do arise after cell division and that there is gradual remodeling of the lamina during interphase. Both of these factors can contribute to the recovery of lamina structure in fibroblasts after constricted migration.

### Both priming of BJ-5ta cells for migration and constricted migration cause 3D genome compartment profile changes

Having explored alterations to and recovery of nucleus morphology, we next asked whether the 3D genome of these cells also experiences alterations and if those alterations were reversible. Since we observed that recovery from the nucleus stress of migration occurs over time after migration, we needed to develop an approach to capture enough cells for Hi-C experiments immediately after migration. We used 3D printing and large 5µm pore mesh to fabricate a 10 cm Transwell insert that would allow collection of larger numbers of cells directly following migration (see Methods for details) **(Sup. Figure 3)**.

We analyzed 3D genomic alterations that occurred in the process of BJ5-ta migration at the compartment scale. In all analyses, we consider both whether the compartment identity “switched” (a change from negative to positive or positive to negative PC1 values at a given genomic location) and also the degree of “shift” toward the A or B compartment (how much the PC1 value increased or decreased, regardless of sign change). We first asked whether the priming of cells by starvation and subsequent spiking with FGF and nutrients but without migration led to any changes in the 3D genome. When comparing S/S to UU, we found that more regions of the genome experienced a shift or switch towards the open, euchromatic A compartment than the closed, heterochromatic B compartment **(Figure 4A)**. Overall, we observed that 8.47% of the genome experienced a shift or switch towards the A compartment with over half of those being a compartment strength shift rather than a compartment switch. This suggests that the preparation for migration caused by starvation and then chemoattractant spiking tends to lessen chromatin compaction. In contrast, only 3.26% of the genome experienced a shift or switch towards the B compartment, and most of those changes were compartment switches without a strong shift in compartment strength. Examples of both shifts and switches are shown in **Figure 4B**. The genomic regions that shift to the A compartment, which would tend to make genes available for activation, contain genes involved in biological processes such as growth factor and stress response, as would be expected for this response to starvation stress followed by FGF exposure **(Figure 4C, full gene lists in Data S1)**.

**Figure 4.**
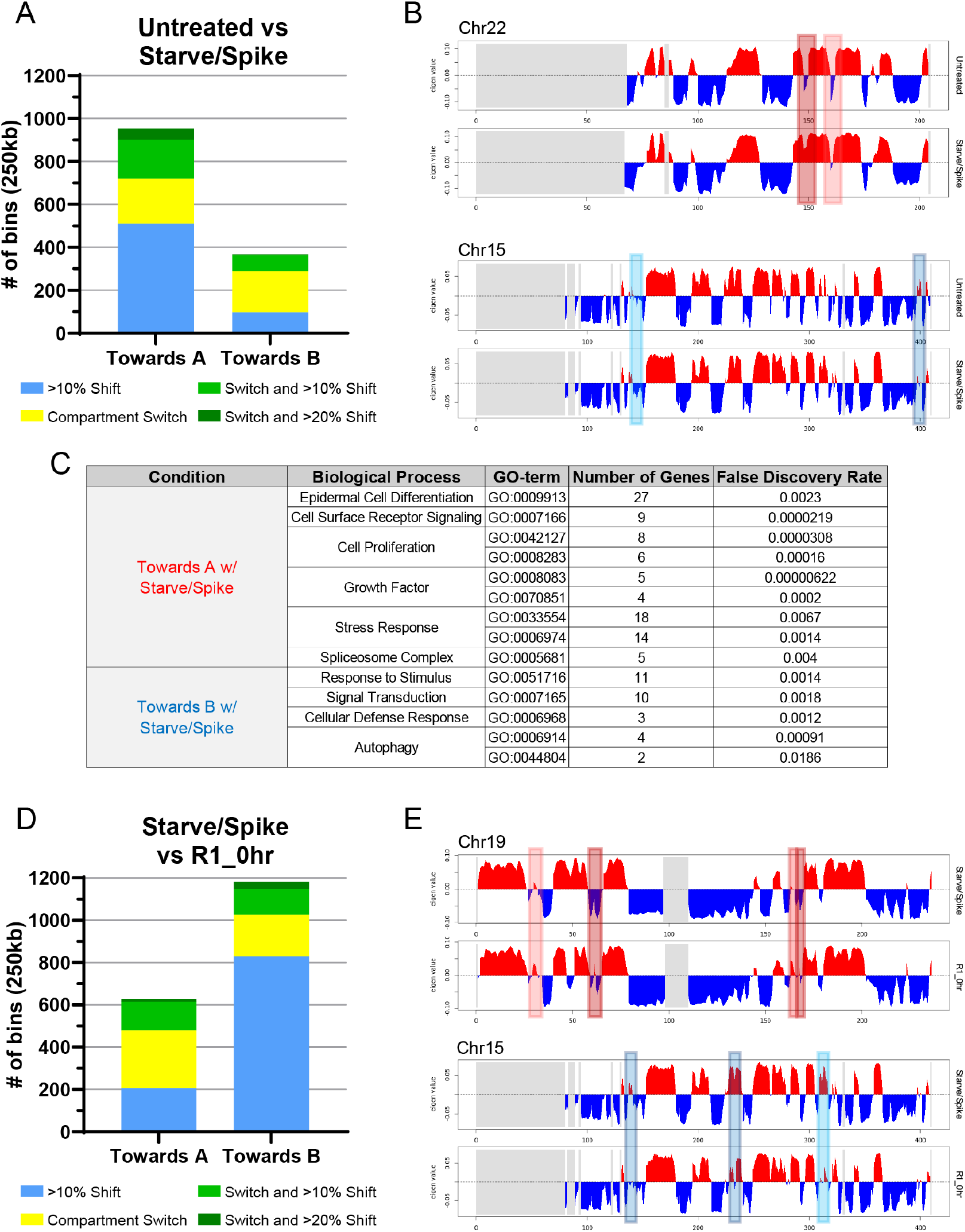
Priming and migration of BJ-5ta cells cause 3D genome alterations at the compartment level. **A)** Comparing the Untreated and Starve/Spike conditions, Venn diagrams show numbers of 250kb bins that switched B to A (dark red) or A to B (dark blue), shifted compartment strength >10% towards A (light red) or towards B (light blue) or experienced both a switch and substantial shift (red or blue intersections). Switched or strength changes to B compartment shown below in Blue. **B)** Examples of bins found in **4A** that switched B to A (dark red boxes), had a strength change >10% towards A (light red boxes), switched A to B (dark blue boxes), or strength change >10% towards B (light blue boxes) **C)** Table of biological processes of genes found in regions that shift towards A or B compartments following starvation and nutrient spiking. **D)** Same as **4A**, but bins are ones that fit the criteria when comparing Starve/Spike to R1_0hr. **E)** Same as **4B** in relation to bins found in **4D**.

We next compared S/S to R1_0hr to determine if passage through constricted migration induced 3D genomic alterations on top of the changes found from starvation and spiking alone **(Figure 4D)**. Interestingly, we observed that 16.09% of the entire genome was changed following constricted migration. Unlike what was seen following starvation and spiking, most of the genomic alterations occurred in bins that shifted or switched towards the B compartment. Most of those alterations were strong shifts rather than switches suggesting that constricted migration leads to chromatin compaction in these cells. For the genomic regions that changed towards the A compartment, almost half of them changed compartment but did not include a strong change in compartment strength, suggesting a more minor shift in spatial organization of these regions. The large number of bins shifting to or towards the B compartment with constricted migration may relate to the previously observed phenomenon that chromatin compaction can facilitate constricted migration (Bustin, 2010) or could relate to the creation of new lamin tethers as internal genomic regions are brought into proximity with the nuclear lamina during constriction. Examples of both switches and shifts that occur following migration can be seen in **Figure 4E**.

### Many migration-associated compartment changes revert to unmigrated state after sequential migration and cell proliferation

Constricted migration did in fact cause 3D genome alterations at the compartment level, so we next looked to determine if cells recovered from these alterations as they did from nucleus deformations over the course of subsequent rounds of migration and cell proliferation. To test this, we used 2 conditions in which cells experienced constricted migration and then were allowed to proliferate for a certain amount of time. A direct comparison to R1_0hr was the R1_96hr which experienced 4 days of growth following migration. In the second condition, R5, cells were migrated through 5 rounds of constrictions but had days to proliferate between each round and after the last round before Hi-C analysis. We compared the difference between each condition, using the S/S pre-migration state as a starting point. We performed hierarchical clustering on all genomic bins that shifted compartments by at least 10% from S/S to R1_0hr to evaluate patterns of recovery or stability in these compartment changes. **(Figure 5A)**. We found that bins that shifted towards the A compartment directly following migration fell into two groups: some shift back towards their unmigrated state (green box), while others remain stable and do not recover (grey box). When looking at R1_96hr and R5, we see that bins within the green box show a lessening difference compared to S/S and some bins even recovering back to UU levels. When looking at the bins that shifted towards B with migration, we saw a larger subset that experienced recovery back to or even beyond that of unmigrated conditions (green box). We also saw a set of bins that do not recover as much and still remain somewhat shifted toward the B compartment after migration and growth (grey box). Examples of bins that changed after migration and recovered after proliferation are shown in **Figure 5B** and **Sup. Figure 4A-D** highlighted by the green boxes. These examples show changes in both directions (towards A and B) that recover back to unmigrated levels in either R1_96hr or R5. **Figure 5C** and **Sup. Figure 4C-D** show examples of bins that change following migration but show no recovery over time (grey highlights).

**Figure 5.**
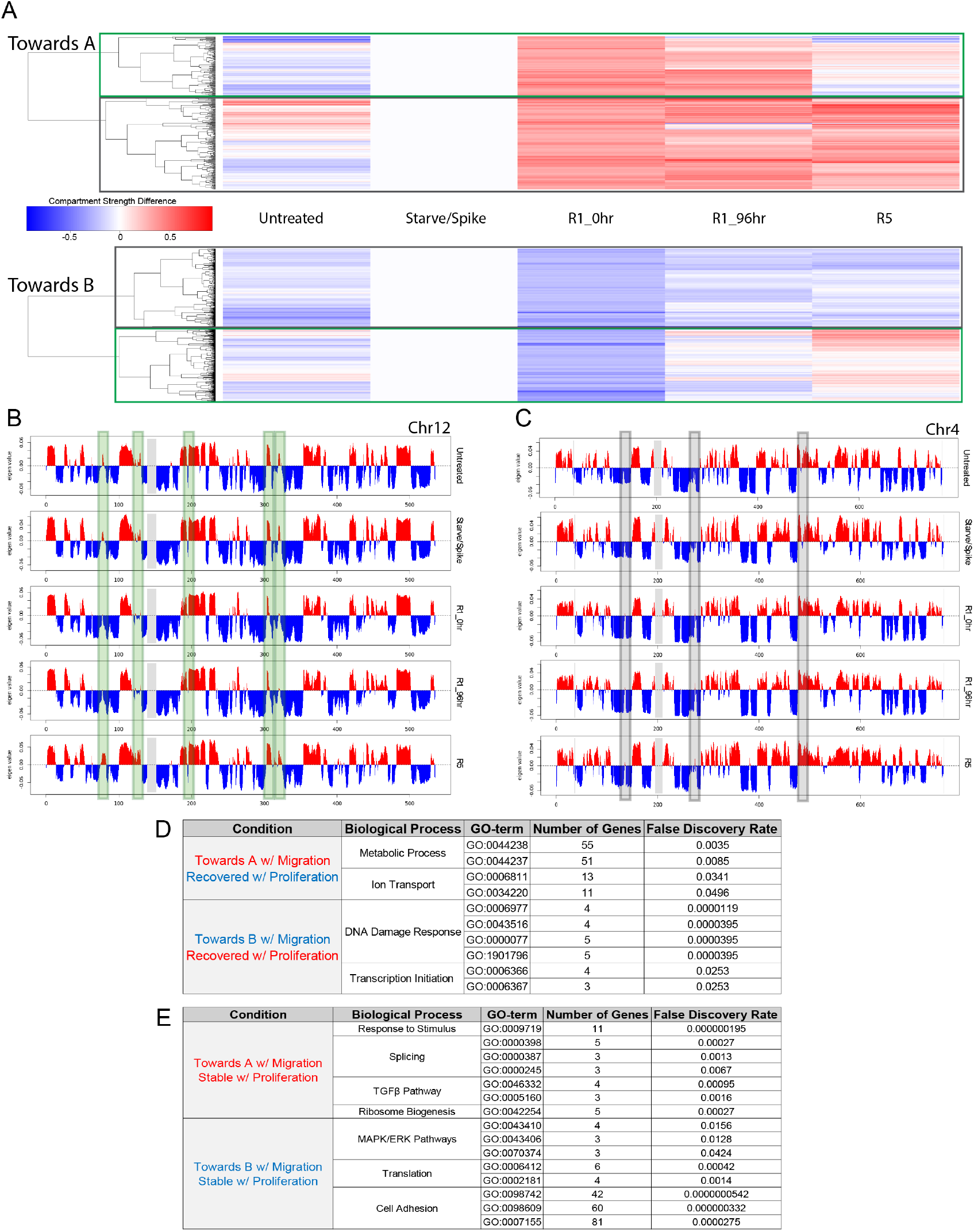
Compartment changes following constricted migration are recovered from over time for some genomic loci. **A)** Hierarchical clustering of all bins that had a greater than 10% shift towards A (top) or B (bottom) when comparing Starve/Spike to R1_0hr. These include bins that switched compartments and had a >10% change as well. Green box highlights bins that show recovery of compartment profile over time, whereas bins that do not recover are highlighted in the grey box. **B)** Examples of bins that exhibit a compartment change following migration that is recovered over time (green boxes). **C)** Examples of bins that exhibit a compartment change following migration but do not recover over time (grey boxes). **D)** Table of biological processes of genes located in regions that changed following migration but returned to unmigrated levels following proliferation (recovered). **E)** Table of biological processes of genes located in regions that changed following migration and remained changed following proliferation (not recovered).

Genes in bins that shifted compartments and then recovered are consistent with a short term response to the stresses of migration. DNA damage response and transcription initiation genes are temporarily shifted toward the B compartment by migration (**Figure 5D, full gene lists in Data S1)**. Meanwhile, genes in processes such as ion transport are transiently shifted toward the A compartment. Ion channels have previously been shown to be important in constricted cell migration and differentially regulated in response to nucleus compression (Mistriotis *et al*., 2024; McCreery *et al*., 2025). Some genes in regions with stable compartment switches are involved in some fundamental processes such as translation, ribosome biogenesis, and RNA splicing (**Figure 5E**). Cell adhesion and cell-cell adhesion genes were enriched in regions that stably shifted toward the B compartment, which could indicate a tendency to downregulate adhesion in favor of migration. We also found enrichment of genes in the MAPK/ERK signaling pathways stably shifted toward B. These pathways have been implicated in altering the cellular mechanical properties and cytoskeletal organization in ways that impact constricted cell migration (Rudzka *et al*., 2019; Samson *et al*., 2022). Conversely, key members of the TGFbeta pathway, including TGFB1, TGFBR3, and SMAD2 were stably shifted to the A compartment. These genes are often involved in the conversion of fibroblasts to myofibroblasts and fibrosis (Chen *et al*., 2025). We see no detectable increase in smooth muscle alpha actin expression after constricted migration **(Sup. Figure 4E)**. Previous reports indicate that these hTERT immortalized fibroblasts rarely convert to myofibroblasts (Harada *et al*., 2021), but we note that the alphaSMA expression after migration was even substantially less than after cell exposure to TGFbeta **(Sup. Figure 4E)**. However, the stable shifting of these genes toward the A compartment after constricted migration suggests increased potential for myofibroblast-like signaling.

In addition to looking at compartment switching, we examined alterations in the frequency of interactions within and between compartments using saddle plots (**Sup. Figure 5)**. Consistent with the prevailing directions of compartment switches, B-B interactions were weakened relative to A-A compartment interactions after starvation and spiking, but then B compartment interactions were strengthened immediately after one round of migration. Compartment interaction strength of both A and B compartments recovered to its pre-migration state after 5 rounds of migration and/or rounds of cell division.

## Discussion

The field of nucleus mechanobiology has revealed that physical forces and deformations can impact the nucleus, chromatin state, and cell phenotype of cell types ranging from cancer to stem cells to immune cells and fibroblasts (Cho *et al*., 2017; Bertillot *et al*., 2022; Kalukula *et al*., 2022; De Corato and Gomez-Benito, 2024). Cancer cells can undergo an epithelial to mesenchymal transition resulting in stably increased migratory capacity after sequential constricted migration (Irianto *et al*., 2017; Golloshi *et al*., 2022; Playter *et al*., 2025). Non-cancerous fibroblasts can likewise experience phenotype shifts as a result of external forces: the persistent tension caused by extracellular stiffness can result in fibroblast conversion to myofibroblasts (Walker *et al*., 2021). Here, we observed that human BJ-5ta fibroblasts repeatedly migrated through nucleus-deforming constrictions did not show the increasing migratory capacity or stable nucleus shape alterations that have been seen in cancer cells. However, these fibroblasts did experience changes to nucleus shape and 3D genome compartmentalization immediately after passage through the constriction.

We observed that the priming of these fibroblasts for migration by starvation and then growth factor exposure initiates changes in nucleus morphology and 3D chromosome compartmentalization before constriction even occurs. These changes are in line with the known nucleus movement and re-orientation and cytoskeletal reorganization that occurs as fibroblasts prepare to migrate (Chang *et al*., 2013). The nucleus elongation and wrinkling we observed both prior to migration and during movement on a 2D surface after migration, are consistent with forces on the nucleus by actin cables or the focal adhesion-anchored actin cap that connects to the nucleus through LINC complexes (Luxton *et al*., 2010; Zhu *et al*., 2018).

Shortly after passage through the constriction, we observed 3D chromosome changes and nucleus deformations that reflect the effects of the constriction on the nucleus and chromatin. Both our compartment switch and compartment interaction strength analyses indicated an increase in B compartment heterochromatin interactions immediately following one round of constricted migration. This observation is in line with both experimental measurements and theoretical models of nucleus compression, in which compressive forces on the nucleus result in chromatin condensation (Damodaran *et al*., 2018; De Corato and Gomez-Benito, 2024). As observed in experiments with transient compression, the increased B compartment interactions after migration were reversible and returned to unmigrated levels after cell proliferation.

Notably, this increase in B compartment interactions after constricted migration is opposite from what was observed after sequential melanoma and breast cancer cell migration (Golloshi *et al*., 2022) or after neutrophil constricted migration (Jacobson *et al*., 2018). We hypothesize that the fibroblast migration more closely reflects what would be expected as a direct but reversible physical consequence of the migration. In cancer, on the other hand, the sequential constricted migration induces an epithelial to mesenchymal transition, and so the compartment strength changes observed likely reflect the alterations to 3D genome organization that facilitate this phenotype switch rather than only physical responses to migration. The difference between the behavior of heterochromatin in fibroblasts and neutrophils may reflect the lack of LaminA/C, a key heterochromatin interactor, in neutrophils.

We found that many of the nucleus shape changes and 3D genome changes are reversible after a recovery period of several days in which cells are able to divide and proliferate. We observed that nuclear lamina remodels both after each cell division and that its structure can also fluctuate throughout cellular movement on a 2D surface during interphase. The chromosome compartment changes which were transient included not only the overall shift toward increased B compartment interactions, but temporary shifts of specific genomic regions which may facilitate gene expression changes needed during or shortly after constricted migration. These include chromatin opening shifts of genes in pathways such as stimulus response and ion transport and shifts toward heterochromatin of transcription and DNA damage response genes. We hypothesize that these temporary shifts facilitate responses such as the previously documented altered regulation of ion channels in response to nucleus compression (Mistriotis *et al*., 2024; McCreery *et al*., 2025).

While wrinkles in the nuclear lamina and a set of spatial compartment changes revert to their original state over time after constriction, we observed other changes which are more persistent and have potential to influence the phenotype of these fibroblasts longer-term. One of the first hints of longer-term adaptive change that we observed was in the seemingly counter-intuitive reduction in severe nucleus deformations after sequential passages through constrictions. The most dramatic lamin wrinkles and deformations were seen after the first round of constriction and lessened in subsequent rounds. We hypothesize that this represents an adaptive response of the cells to enable future migration with less long-term disruption to nucleus geometry. This ability to recover is itself a notable phenotypic change that seems to emerge as a result of the first rounds of constriction. This change may be facilitated by some of the compartment changes that do not recover after constriction but instead are stable through rounds of cell division in culture and over the course of multiple rounds of constriction. We found that these more stable compartment changes included shifts toward the B compartment for genes in pathways such as MAPK/ERK signaling and cell-cell adhesion, factors known to be involved in controlling cell migration behavior (Rudzka *et al*., 2019; Samson *et al*., 2022). Shifting cadherin proteins such as CDH2, 9, 12, and 18 and claudin tight junction proteins (CLDN22-24) toward the B compartment long term could favor individual cell migration over collective migration or cell-cell adhesions (Kaflak-Hachulska *et al*., 2003). Notably, TGFbeta and other proteins associated with TGFbeta signaling were among those stably shifted into the A compartment after constricted migration. While we did not observe associated smooth muscle alpha actin expression that would indicate such TGFbeta signals were triggering myofibroblast formation, these compartment shifts could indicate that these genes are poised for future activation and could make myofibroblast formation more likely after constricted migration. Also, we note a caveat here about our use of hTERT transformed fibroblasts--while enabling persistent long-term culture of these fibroblasts over extended migration experiments, the TERT expression may suppress myofibroblast conversion that might otherwise be triggered by the physical stress of migration (Harada *et al*., 2021). Therefore, additional future work will be needed to validate the results seen here in primary human fibroblasts.

Overall, our results suggest that fibroblasts share some similarities with cancer cells in the nuclear effects of constricted migration, but the phenotypic impact and stability of those changes is quite different. The fact that phenotypic and genome structure changes in cancer cells are more dramatic after constricted migration likely reflects the increased heterogeneity and chromosome structure plasticity observed in cancer cells that have already lost their original cell identity (Roy and Hebrok, 2015; Frederick *et al*., 2025) compared to the terminally differentiated fibroblast cell type. Our results shed light on how fibroblasts, as “professional” migrators adapt and respond to the forces of constriction during processes such as wound healing, recovering from these stresses to be able to respond again to a new stimulus in the future.

## Methods

### Cell Culture

BJ-5ta Cells were purchased from ATCC (CRL-4001). They were grown in a 4:1 ratio of DMEM (Corning, 10-013-CV) and 1x Medium 199 (Gibco, 11-150-059), supplemented with 10% FBS (Corning, 35-010-CV), 0.01 mg/mL hygromycin B (Corning, 30-240-CR), 1% Pen-Strep (Gibco, 14140-122), and 1% L-Glutamate (Gibco, 25030-081).

### Transwell Assays (24-well Plate)

Transwell filters with 5µm pores (VWR-10769-236) were used. The bottom of each filter was coated with 40µL of 10ug/µL fibronectin (Corning, 354008) for ∼30 min. 100,000 BJ-5ta cells in 100µL of fully supplemented media were placed into the top well of the Transwell with 500µL of fully supplemented media in the bottom chamber. Cells were allowed to attach overnight. The following day both the top and bottom chamber’s media were replaced with a 4:1 ratio of DMEM and 1x Medium 199, but with 1% BSA instead of 10% FBS. Cells were starved overnight. The following day, the bottom chamber’s media was replaced with a 4:1 ratio of DMEM and Medium 199 supplemented with 20% FBS and 5 μg/mL of FGF (Gibco, PHG0368V).

After the 24hr incubation, migration efficiency was quantified as follows; first, freely floating cells were removed from the top of the filter (unmigrated cells) and from the well beneath the filter (migrated cells) and placed in two separate tubes. Then, 400μL of trypsin was added into the bottom chamber of the 24-well plates, and 200μL of trypsin was added into the top chamber to detach any remaining attached cells. Recovered cells after trypsinization were added to the unmigrated or migrated tubes, accordingly. Cells were spun down (1,000 rpm, 5 min) and resuspended in 100µL of DMEM. 15µL from each tube was collected and counted (using trypan blue) on a hemacytometer to calculate % migration such as: #bottom/(#top+#bottom).

For sequential rounds of migration, the migrated cells were seeded into a new well of a 24-well plate to expand. When cells reached 80–90% confluency, another Transwell migration was performed as previously described. Only the bottom cells were collected and continued to be used for sequential rounds of migration.

### Immunofluorescent Confocal Imaging

Approximately 50,000 BJ-5ta cells for each condition were seeded into 35-mm glass bottom dishes (MatTek, P35GC-1.5-10-C). Cells were allowed to attach overnight and then cross-linked with 4% formaldehyde for 10 min followed by three, 5-min washes with PBS. After washing, cells were permeabilized with permeabilization buffer (10% goat serum, 0.5% Triton in PBS) for 1 h at room temperature. After the incubation, primary antibody for LaminA/C (Santa Cruz, sc-376248), diluted in antibody dilution buffer (5% Goat serum, 0.25% Triton in PBS), was added and incubated overnight at 4°C. After primary antibody incubation, cells were washed three times with PBS for 5 min for each. Cells were then incubated with secondary antibodies (Alexa Fluor 488, Invitrogen, R37120) per manufacturer’s directions (two drops of secondary antibody/1 mL of PBS) for 30 min at room temperature. After secondary antibody incubation, cells were washed three times with PBS and sealed using mounting media with DAPI (Invitrogen, P36962). Slides were treated in the mounting media for 24hr before imaging. Cells were imaged using a Leica Sp8 Confocal microscope equipped with a 63x oil immersion objective. For αSMA experiments, cells were seeded onto 24 well plate coverslips (Fisher Scientific, 50-194-4702) with either no treatment, starvation and FGF spike-in as described earlier, directly after 5um Transwell migration, or after migration and allowed to recover for 48hrs. Cells were then fixed as above and stained with an αSMA primary antibody (Invitrogen, MA1-06110) overnight in 4C. Secondary antibody and washes were done as above. Cells were then imaged on a Leica Sp8 Confocal microscope with a 40x water immersion objective. For a positive control for αSMA induction, BJ-5ta cells were treated with 5ng/mL TGFbeta for 48hrs before fixing and staining as described above on 24 well plate sized coverslips.

### Nucleus Characteristic Analysis

Aspect Ratio and Solidity measurement were completed using the Shape Descriptors plugin in Fiji based on region of interest overlays created from the LaminA/C signal only. Standard Deviation measurements were made using the standard deviation tool in Fiji on the same regions of interest overlays made from the LaminA/C signals. The cumulative plot for LaminA/C distribution was created in R studio using cumulative plot command on the standard deviation results from Fiji. EFC ratios were determined using the MATLAB script described in Tamashunas et al (Tamashunas *et al*., 2020b) on .tiff images of the LaminA/C signal only for the same nuclei as the shape descriptors.

### LaminA/C-GFP Transduction

BJ-5ta cells were stably transduced with N-terminal tagged GFP-LaminA using the pBABE-puro-GFP-wt-lamin A plasmid (Plasmid #17662, Addgene). The plasmid was packaged and delivered using the Phoenix-AMPHO (ATCC® CRL-3213) system. After viral supernatants were filtered and added to BJ-5ta cells, the cells expressing the construct were selected with puromycin.

### FACS Sorting/Flow Cytometry

BJ-5ta GFP-LMNA cells were analyzed on a Cytek Northern Lights cytometer to determine relative levels of GFP signal per cell before cell sorting. An Unstained population of cells was also used as a negative control to set a threshold value for GFP-LMNA expression. Cell sorting was completed on a Sony 9000 cell sorter where collection of only lowly positive GFP cells occurred. Anything negative for GFP or above our threshold was removed. An Unstained population of cells was also used as a negative control to set a threshold value for GFP-LMNA expression.

### Live Imaging Movies and Analysis

20,000 GFP-LMNA sorted BJ-5ta cells that had been passed through 1 round of constricted migration (as described above) were seeded into a fibronectin coated (10µg/µL) Zeiss eight-chambered slide and allowed to attach. Following attachment, cells were taken to either a Zeiss Confocal (63x) or a Keyence microscope (20x) and z-stack images were taken every 10 minutes for either 8 hours or 72 hours respectively. Analysis of movies followed the same principles as the nucleus characteristic analysis as described above.

### 10cm Transwell Creation

CAD was used to create the insert (see **Data S2** for .stl file), which was printed on a FormLabs Form 3B resin printer out of FormLabs Clear V4 resin. The inserts were then processed in FormLabs wash and cure stations to remove any excess resin and ensure a complete cure. Supports were then removed, and the final parts lightly sanded to ensure a smooth, flat surface on which to glue the migration membrane.

### 10cm Transwell Assay

A 5µm pore mesh (Sterlitech, PCT5014220) was attached to the bottom of the insert using a 2-part epoxy spread on by a P1000 pipette tip. Epoxy was dried for 24hours at room temperature before trimming any excess mesh on the outside of the well. Completed inserts were sprayed with 70% ethanol and allowed to dry for ∼2 hours for sterilization. The sterile and completed Transwell inserts could then be placed in any 10cm dish bottom for migration assays. To coat the bottom of the filters in fibronectin, 1mL of 10µg/µL fibronectin was placed on parafilm and filters placed on top for ∼30 mins with any excess being removed prior to migration assays. Migration experiments followed the same protocol as the 24-well plates above with adjusted cell number added (5 million), and volumes for top and bottom chambers being 7mL and 9mL respectively. For incubation steps, the entire apparatus was placed inside a 15cm dish for sterility purposes. Cells were still detached with trypsin and either went through fixation and nucleus isolation for Hi-C (R1_0hr) or seeded into T75 flasks and allowed to grow for 4 days before fixation and nucleus isolation for Hi-C (R1_96hr).

### Nucleus Isolation

Nucleus isolation protocol was adapted from Moniot-Perron et al (Moniot-Perron *et al*., 2023). In short, BJ-5ta cells were isolated in 500µL of cold PBS with 10% FBS and pushed through a 22µm cell strainer to avoid clumps. They were then fixed in 10mL of a 2% formaldehyde solution for 10 minutes at RT on a shaking platform. Crosslinking was quenched with 1.425mL of 1M Glycine and placed on ice for 5 minutes before being spun down at 225g for 8 minutes in a 4C centrifuge. Supernatant was removed and cells were suspended in 5mL of cold Lysis buffer (600µL 1M Tris-HCl, 360µL 5M NaCl, 120µL 0.5M EDTA, 300µL 20% NP-40, 600µL 20% Triton, 240µL Complete Protease Inhibitors, 9.8mL of MilliQ Water). Cells were incubated on ice for 10mins before spinning down for 5min at 400g. Supernatant removed and isolated nuclei were flash frozen in liquid nitrogen and stored at – 80C.

### Hi-C Experiments and Analysis

All Hi-C experiments were performed on isolated nuclei (described above) using the Arima-Hi-C+ Kit from Arima Genomics following the protocol for Mammalian Cell Lines (A160134 v01) and preparing the libraries using Arima recommendations for the NEBNext Ultra II DNA library prep kit (protocol version A16041v01). Sequencing was performed through GeneWiz using an Illumina NovaSeq with 150bp paired end reads. Sequencing reads were mapped to the human genome (hg19), filtered, and iteratively corrected by Hi-CPro (https://github.com/nservant/Hi-C-Pro). For library quality and mapping statistics, see **Sup. Table 1**.

Compartment analysis was performed by running principal component analysis using matrix2compartment.pl script in the cworld-dekker pipeline available on GitHub (https://github.com/dekkerlab/cworld-dekker). The PC1 value was then used to determine compartment identity for 250kb binned matrices. We considered PC1 values greater than 0.01 to be A compartment and less than −0.01 to be B compartment. To define compartment switches between conditions, PC1 values must go from positive to negative (A to B) or negative to positive (B to A). For compartment shifts between conditions, the difference between conditions had to have a PC1 value change greater than 0.02 (Towards A) or less than -0.02 (Towards B). This amount of PC1 change correlates with a 10% change in compartment score. For a 20% change in compartment score, PC1 values must have changed more than 0.04 or less than –0.04.

Compartment differences and clustering for changes following migration were completed as follows: All bins that experienced a >10% eigen vector value change (+/-0.02) between S/S and R1_0hr were collected. Eigen vector values for all conditions on those selected bins were ordered as follows: UU, S/S, R1_0hr, R1_96hr, R5. For normalization to S/S, the difference between all eigen vector values and that of S/S were calculated (S/S difference was 0 for every bin with shifts towards A being positive values and shifts towards B being negative values for the other conditions). Then, bins that shifted towards A and towards B in R1_0hr were separated into individual lists for clustering analysis. Clustering analysis was completed in R Studio using all bins shifted (shown in **Figure 5A**). To see how genomic locations respond to priming, we started with the list of bins that shifted when comparing UU to S/S and followed the same process as before. We then used UU eigen vector values in place of S/S to calculate differences for clustering (UU difference being 0 for each bin).

### Gene Function Enrichment Analyses

We first identified all genomic bins that switched compartment and had at least a 10% shift in PC1 value between conditions of interest. Next, UCSC tables were used to extract lists of genes in these genomic bins. STRING database analysis (Szklarczyk *et al*., 2023) was then used to search for biological function enrichment of these gene lists using the following approach: 1) Interaction networks were defined using Textmining, Experiments, Databases and Co-expression at medium confidence (0.4). 2) Informative functional enrichments in the entire list were first noted. 3) Kmeans clustering was run to identify the number of disconnected graphs in the network, and then functional enrichments were noted within each of these smaller clusters. This was done because often there are multiple meaningful subgroups of gene functions, but none of them will show up as “significant” out of the larger set. 4) Separately, DBScan clusters were generated to enable a search for small functional complexes. This multi-tiered approach enabled us to identify the shared functions of genes that changed compartment in a way that would be missed if only enrichments in the whole gene set were considered. “Informative” functional enrichment means a term that is statistically enriched in a given gene set (FDR < 0.05) but is neither too narrow (for example, 2 genes out of 500 in a certain function) nor too broad (that is, terms like “positive regulation”) to interpret. Functionally related biological terms were grouped together conceptually with each GO term ID annotated in the final tables.

### Saddle Plots

Multiresolution cool files were created from the HiCPro validpair files using the cooler package https://github.com/open2c/cooler. Saddle plots and compartment strength were calculated using the saddle plot functions in coolTools (https://cooltools.readthedocs.io/en/latest/notebooks/compartments_and_saddles.html).(Open2C *et al*., 2024). The bins with the strongest 20% of positive (for A) or negative (for B) PC1 values were used to calculate each interaction category (AA, BB).

## Glossary

Untreated (**UU**): Cells grown in normal media, no migration
Starve/Spike (**S/S**): Cells starved overnight in 1% bovine serum albumin (BSA) then spiked with 5 μg/mL fibroblast growth factor (FGF) and cell culture media containing 20% FBS for 24 hours, but no constricted migration
BottomR1-4 (**R1-4**): Cells were starved and spiked, then allowed to migrate through 5 µm pore Transwell filters 1-4 times
BottomR1 No Growth (**R1_0hr**): Cells were starved and spiked, allowed to migrate through a 10 cm 5 µm pore Transwell, then fixed instantly afterwards for Hi-C.
BottomR1 w/ Growth (**R1_96hr**): Cells were starved and spiked, allowed to migrate through a 10 cm 5 µm pore Transwell, then allowed to proliferate for 96 hours before fixing for Hi-C.
BottomR5 (**R5**): Cells were migrated through 5 sequential rounds of 24-well plate 5 µm pore Transwell filters, given 96hr to proliferate after each migration round and before fixing for Hi-C.

## Acknowledgments

We thank all members of the McCord lab for helpful discussion and feedback. We acknowledge the UTK Advanced Microscopy and Imaging Center and Genomics Core for assistance with confocal microscopy and sequencing.

## Author contributions

C.P., S.J.B, and R.P.M. designed the study. C.P., S.J.B., R.D.M., T.B.S., A.S. and K.P. performed experiments and analyzed data. P.Y. assisted in device creation. C.P. and R.P.M. wrote the manuscript with contributions from all authors.

## Data Availability

Raw and processed Hi-C data files are available at GEO accession number GSEXXXXX. Imaging data are available at Dryad at https://doi.org/10.5061/dryad.mcvdnckff.

