## Supplementary Figures for "Reversibility of Nuclear and 3D Genomic Changes in Non-Cancerous Fibroblasts After Constricted Migration"

**Supplementary Information: 5 figures, 1 table, 6 movies, 2 data files**

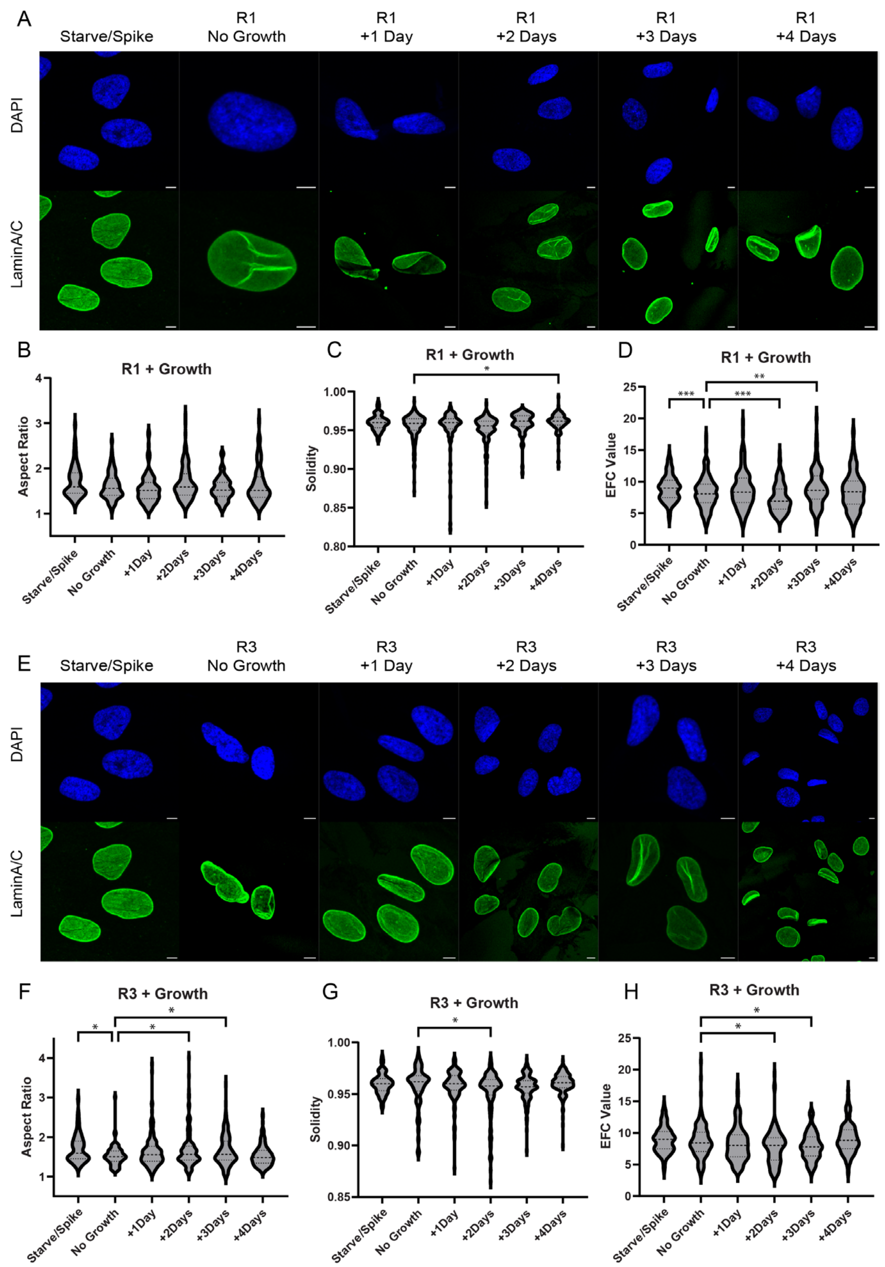


**Sup. Figure 1 – Nucleus deformations recover over time for R1 and R3**

**A)** Confocal images (63x) of BJ-5ta nuclei stained with LaminA/C (green) and DAPI (blue) of S/S and R1 migrated cells immediately after migration (no growth) and all the way through 4 days of proliferation. Scale=10μm. Aspect Ratio **(B),** Solidity **(C)** and EFC Ratio **(D)** measurements for conditions shown in **S1A**. (****P<.0001, one-way ANOVA. N= 68-147 nuclei per condition.) **E)** Confocal images (63x) of BJ-5ta nuclei stained with LaminA/C (green) and DAPI (blue) of S/S and R3 migrated cells immediately after migration (no growth) all the way through 4 days of proliferation. Scale=10μm. Aspect Ratio **(F),** Solidity **(G)** and EFC Ratio **(H)** measurements for conditions shown in **S1E**. (***P<.001, **P<.01, *P<.05, one-way ANOVA. N = 68-177 nuclei per condition.) In all violin plots, thick dashed line indicates median and thin dashes indicate 25$^{th}$ and 75$^{th}$ percentiles, respectively.

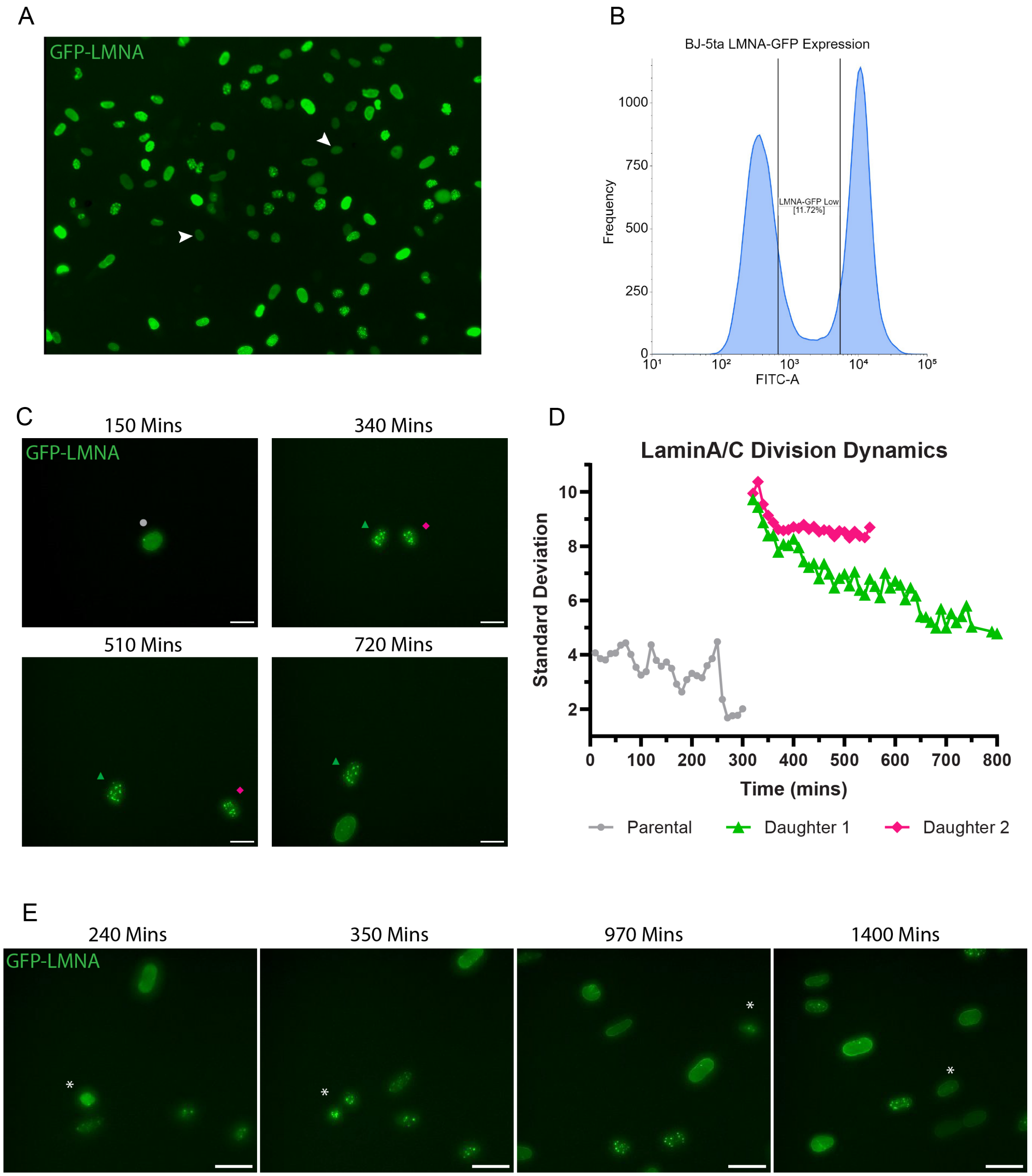


**Sup. Figure 2 – LaminA/C-GFP forms puncta following cell division**

**A)** 10x EVOS image of BJ-5ta cells following transduction of GFP-LMNA. White arrows indicate nuclei that expressed levels of GFP deemed biologically normal and were selected for later on. **B)** Histogram of GFP-LMNA expression used to sort for lowly positive cells (area labeled "GFP-LMNA Low") **C)** Confocal snapshots of **Movie 4** showing a single nucleus before division, right after division, and for a prolonged time after division. Scale = 10μm **D)** Frame by frame measurements of GFP-LMNA standard deviation of the parental and subsequent daughter nuclei shown in **S2C. E)** Confocal snapshots of **Movie 5** showing another example of GFP-LMNA puncta formation following division (dividing nucleus marked by asterisk) that dissipates over time. Scale = 20μm

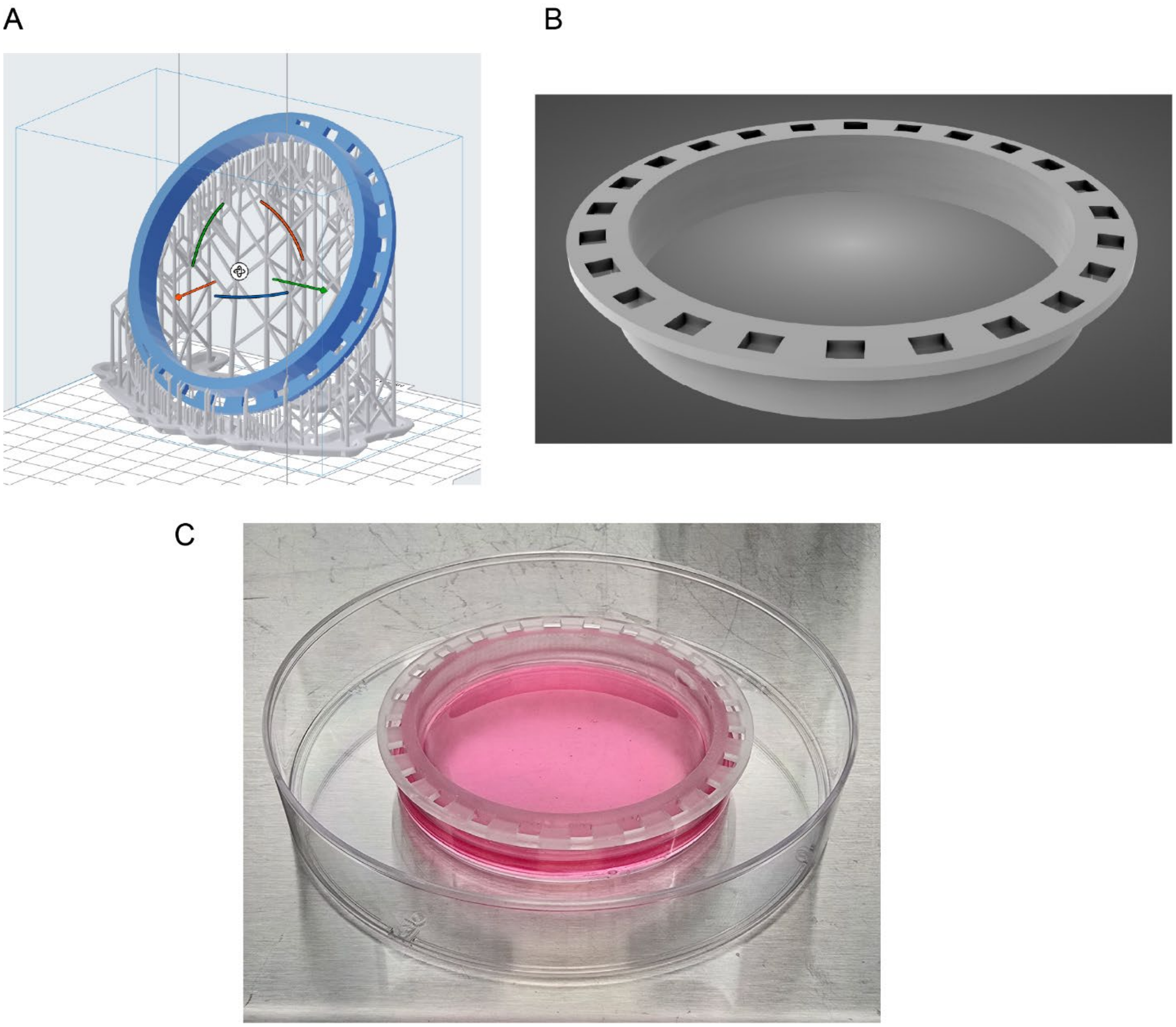


**Sup. Figure 3 – Schematic of 10 cm 5 µm pore Transwell**

**A) I**mage of the 10 cm Transwell insert from the .stl file (see **Data S2**) used for 3D printing. **B)** Top view of Transwell insert without supports. **C)** Completed insert with 5µm pore mesh with top and bottom wells filled with media placed inside a 15cm dish.

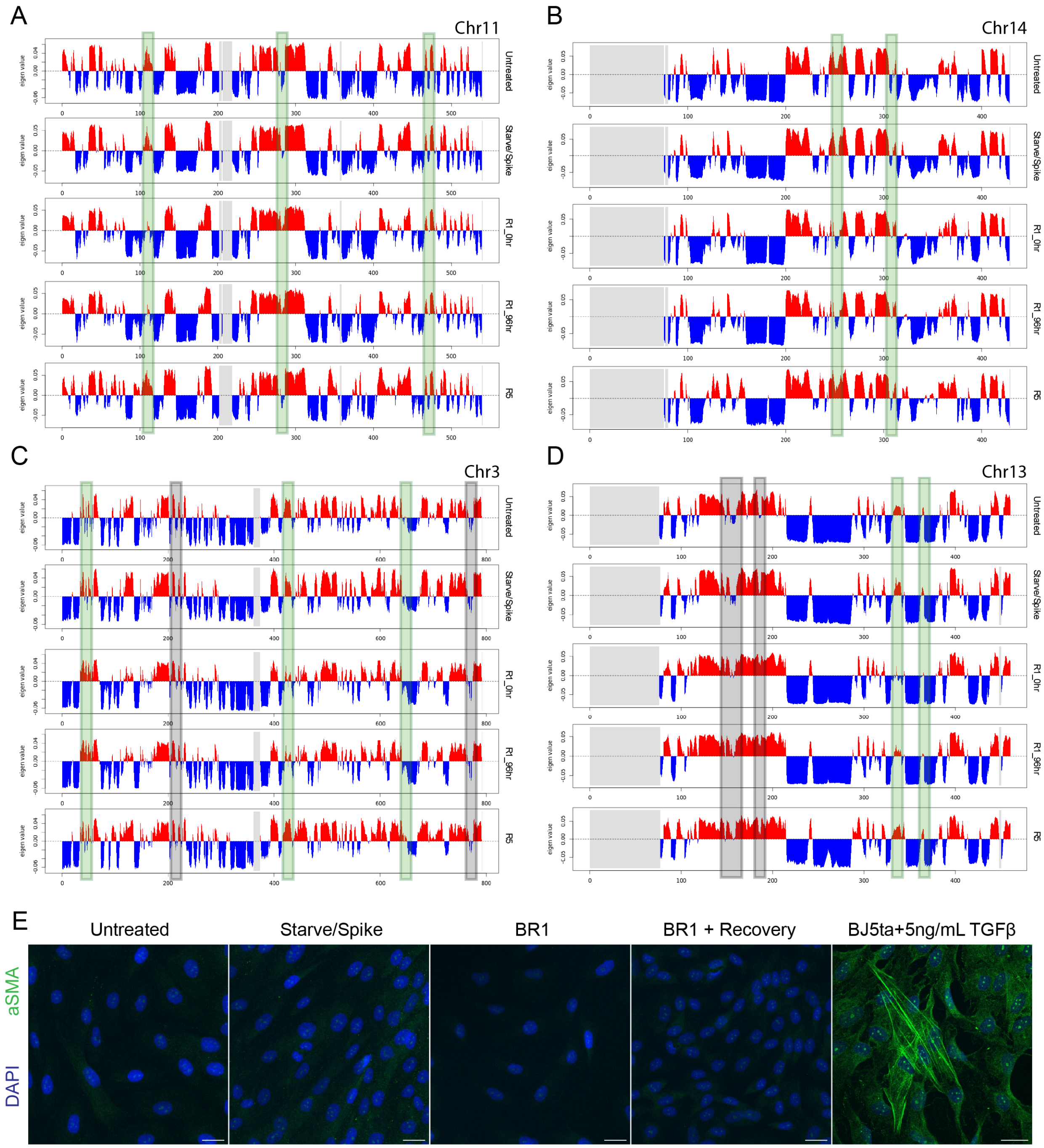


**Sup. Figure 4 – Extra examples of recovered and unrecovered genomic bins following migration with subsequent proliferation and αSMA induction images**

**A-D)** More examples of genomic bins that recover (green box) or do not recover (grey box) from changes caused by migration following proliferation. **E)** Confocal images of BJ-5ta cells before and after migration stained for αSMA expression. Far right panel is 5ng/mL TGFbeta induced BJ-5ta cells used as a positive control for αSMA induction. Scale = 25µm.

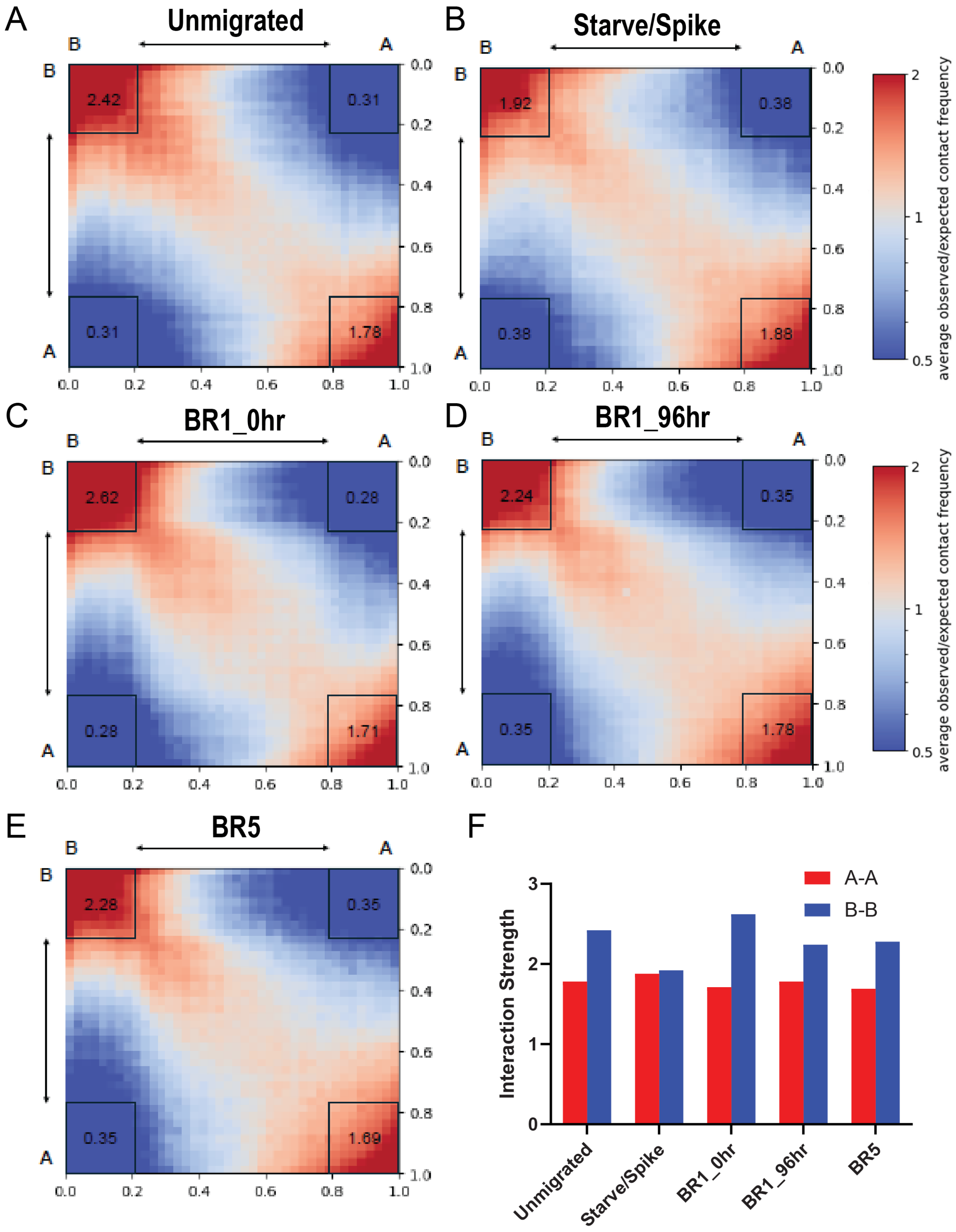


**Sup. Figure 5 – Hi-C Interaction Saddle Plots**

**A-E)** Saddle plots of Untreated, Starve/Spike, BR1_0hr, BR1_96hr, and BR5 showing interaction strengths of B-B, A-A, and A-B interactions for all conditions. **F)** Graphical representation of the interaction strength of the top 20% of bins for A-A and B-B interactions for all conditions.

**Supplementary Table 1.** Hi-C sequencing and mapping statistics.

| Sample | Total Reads | Both Sides Mapped | % Both Sides Mapped | %Dangling Ends | Valid Pairs | Unique Valid Pairs | %Cis | %Cis >20 kb |
|---|---|---|---|---|---|---|---|---|
| BJ5ta-Untreated | 294,966,376 | 184,223,391 | 62.46 | 2.77 | 175,694,557 | 112,085,567 | 87.14 | 57.71 |
| BJ5ta-StarveSpike | 845,064,555 | 466,614,413 | 55.22 | 6.96 | 412,687,750 | 236,424,015 | 87.41 | 55.86 |
| BJ5ta-BR1-0hr | 319,061,522 | 193,557,999 | 60.66 | 8.08 | 169,122,777 | 128,296,113 | 87.4 | 60.2 |
| BJ5ta-BR1-96hr | 735,286,374 | 420,307,301 | 57.16 | 6.18 | 378,191,490 | 246,182,971 | 87.79 | 59.29 |
| BJ5ta-BR5 | 335,394,720 | 204,233,196 | 60.89 | 6.06 | 178,243,875 | 97,625,962 | 89.07 | 46.52 |

**Movie Captions/Legends**

**Movies 1-3 –** BJ-5ta cells transduced with GFP-LMNA migrated through 1 round of 5μm pores. 24hours after seeding, images taken for 8 consecutive hours at 63x on a Zeiss confocal microscope every 10 minutes. Scale = 10μm.

**Movie 4 & 5 –** BJ-5ta cells transduced with GFP-LMNA migrated through 1 round of 5μm pores. 24hours after seeding, images taken for 72 consecutive hours at 20x on a Keyence confocal microscope every 10 minutes. Scale = 20μm.

**Movie 6 –** Unmigrated BJ-5ta cells transduced with GFP-LMNA imaged on a Zeiss confocal at 20x. Images taken for ~16 hours every 10 minutes.

**Data S1 –** Tables of all genomic bin locations and associated genes that shifted or switched compartment in starvation/spiking condition or after 1 round of migration, annotated with whether the compartment status recovered or not after sequential migration and proliferation.

**Data S2 –** File (.stl) describing the parameters for 3D printing the large Transwell device.
